# Robust and Efficient Assessment of Potency (REAP): A Quantitative Tool for Dose-response Curve Estimation

**DOI:** 10.1101/2021.11.20.469388

**Authors:** Shouhao Zhou, Xinyi Liu, Xinying Fang, Vernon M. Chinchilli, Michael Wang, Hong-Gang Wang, Nikolay V Dokholyan, Chan Shen, J Jack Lee

## Abstract

The median-effect equation has been widely used to describe the dose-response relationship and identify compounds that activate or inhibit specific disease targets in contemporary drug discovery. However, the experimental data often contain extreme responses, which may significantly impair the estimation accuracy and impede valid quantitative assessment in the standard estimation procedure. To improve the quantitative estimation of the dose-response relationship, we introduce a novel approach based on robust beta regression. Substantive simulation studies under various scenarios demonstrate solid evidence that the proposed approach consistently provides robust estimation for the median-effect equation, particularly when there are extreme outcome observations. Moreover, simulation studies illustrate that the proposed approach also provides a narrower confidence interval, suggesting a higher power in statistical testing. Finally, to efficiently and conveniently perform common lab data analyses, we develop a freely accessible web-based analytic tool to facilitate the quantitative implementation of the proposed approach for the scientific community.

## Introduction

The median-effect equation is a unified theory in medicine to describe the dose-response relationship and identify agents or their combinations that activate or inhibit specific disease targets^1^. It is a fundamental method established based on the pharmacological principle of mass-action law^2^. As the common link for many biomedical systems, it has been used extensively to analyze *in vitro* experimental data and evaluate the potency of related drugs^3–6^.

In practice, the median-effect equation can be estimated for drug efficacy or pathway inhibition from normalized data generated from experimental studies. Without knowing the true dose-effect curve during the experimental design and data collection, it is common to observe extreme values of (un)affected cell fraction that is close to the response of either 0 or 100% in the analytic dataset. Quantitatively, it poses a special analytic challenge to estimate the median-effect question in practice. The standard estimation approach, often based on a linear regression model after a logit transformation^7,8^, could suffer badly from poor estimation in such situations. **Figure 1** illustrates a preliminary example in that the standard approach is deficient in describing the median effect curve with a perturbation in one extreme data point. The variation in real experimental data, mostly caused by unavoidable measurement error, often at a much larger degree, therefore challenges the reliability of result presentation and interpretation for many drug assessment studies.

**Figure 1.**
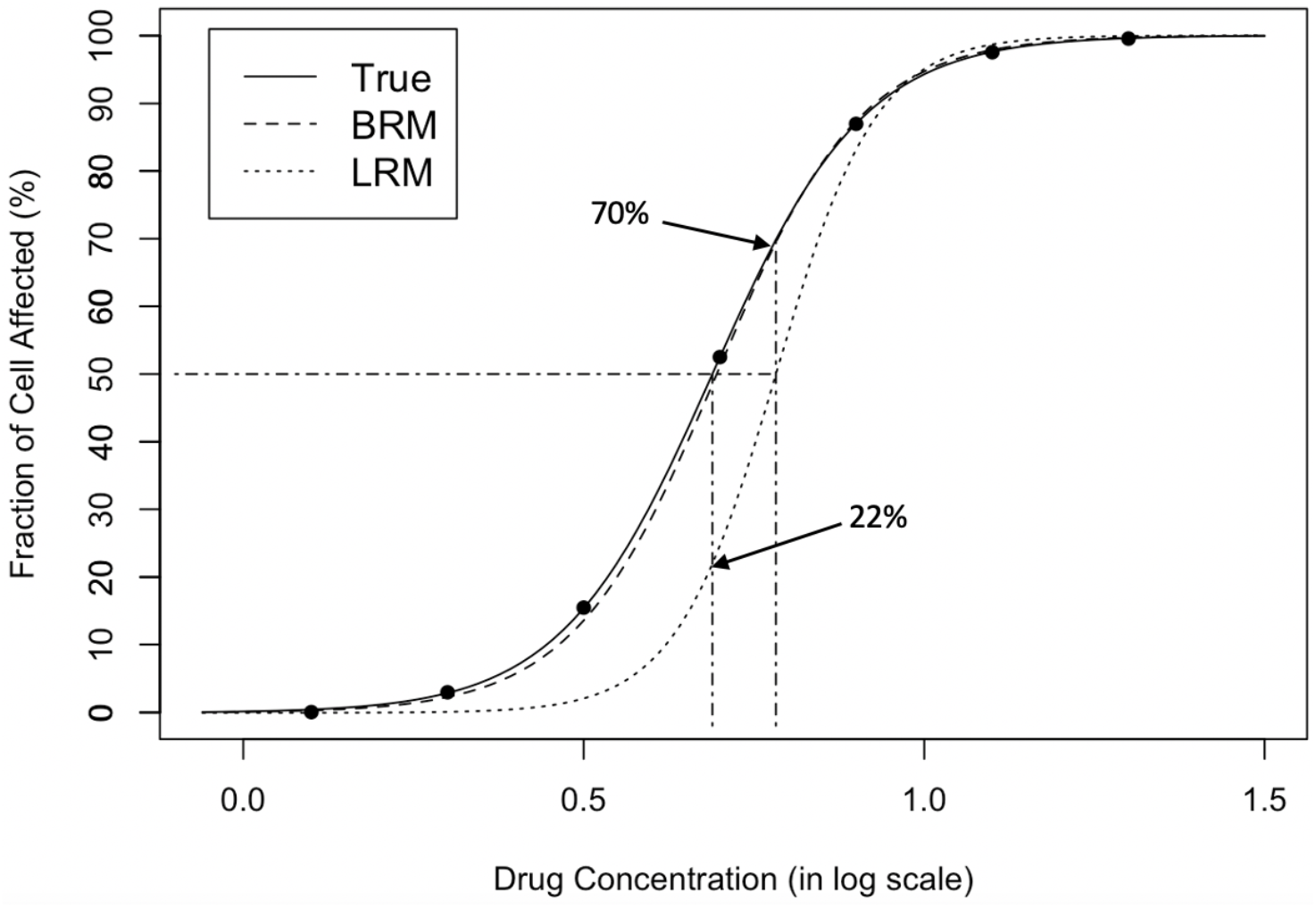
Dose-response curve fitting with extreme observations. The original data points are on the true curve. The first data point (boxed) is changed from 0.005 to 1e-6, referring to a small white noise that cannot be visually recognized. The change leads to the obvious departure between the estimated curve by linear regression model (dotted) and the true curve (solid), which demonstrates that standard regression is sensitive to extreme values. The response at the true IC_50_ (dotdashed, vertical, left) is only 22% from the estimated curve; the estimated IC_50_ (dotdashed, vertical, right) corresponds to the 70% fraction of cell affected, effecting a substantive 20% inflation (50% -> 70%) in estimation error. In contrast, the estimated curve by beta regression model (dashed) is almost overlapped with the true curve (solid), which shows that BRM is much more robust to extreme values. LRM: linear regression model (after logit transformation); BRM: robust beta regression model.

Additionally, the modeling strategy by deleting extreme values may not be feasible in many situations^9^. For example, a meaningful drug concentration could consist of high inhibition (>90%) or low cell viability (<10%) in cancer research. It is not logical to ignore extreme observations when they are indeed biologically relevant for the target effect, not even to mention an associated loss of power and accuracy by leaving fewer data points for estimation. As illustrated in **Figure 2**, deleting the extreme values couldn’t eliminate the estimation bias, but only impaired the efficiency of interval estimation with wider nominal 95% confidence intervals (C.I.) and harmed the estimation accuracy with worse coverage probabilities.

**Figure 2.**
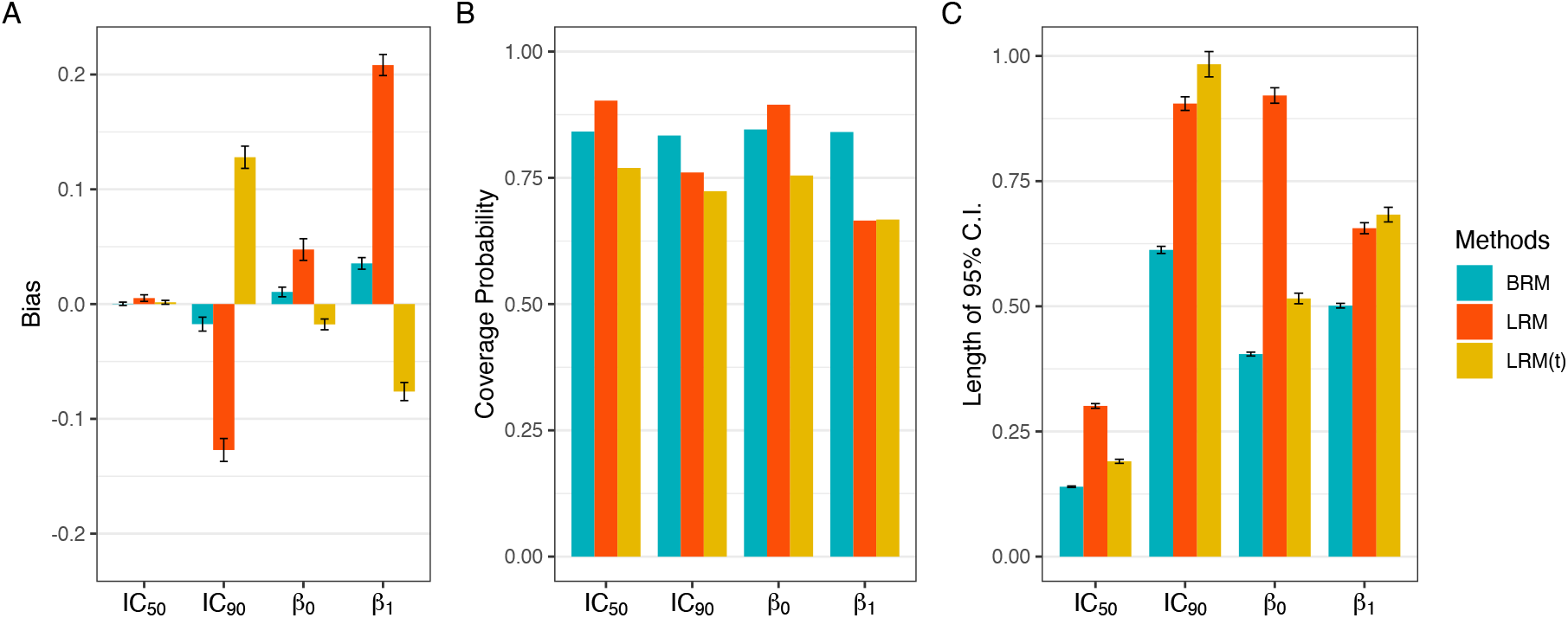
Comparison of estimation efficiency and accuracy using linear regression model and beta regression model. Deleting the extreme values couldn’t eliminate the bias (panel A), but only harmed the accuracy with worse coverage probabilities (panel B) and impaired the efficiency of interval estimation with wider nominal 95% confidence intervals (panel C). 1000 data sets were generated following the data simulating process described in Supplementary Notes, using the dose sets and true dose-response curve under 7 dose setting with a precision parameter of 100. Responses ⩽ 5% or ⩾ 95% were considered extreme responses. BRM: beta regression with extreme data points; LRM: linear regression model (after logit transformation) with extreme data points; LRM(t): linear regression model (after logit transformation) with truncated dataset after deleting extreme values.

Furthermore, it is dubious to apply the constant error variance, a default assumption in standard linear regression modeling, in dose-response estimation. As an assumption can be examined with repeated measures, many dose-response data have indicated either a constant variance before logit transformation or a positive correlation with drug dose. It is incongruous to apply linear regression if the assumption is violated due to error heteroscedasticity^10,11^. Therefore, it is essential to develop a robust quantitative approach to estimating the median-effect equation.

Here, we introduce a novel approach to improving the quantitative assessment of dose-response relationship and drug potency, together with a user-friendly web-based analytic tool to facilitate the implementation. The proposed method to estimate the median-effect equation is established in the robust beta regression framework, which not only takes the beta law to account for non-normality and heteroskedasticity^12^, but also minimizes the average density power divergence (DPD) using a tuning parameter, which compensates for the lack of robustness against outliers under the standard beta regression^13^. Results from simulation studies under various scenarios confirm that the proposed approach consistently gives robust estimation for the median-effect equation. Particularly, we examine two important measures for drug binding affinity: the Hill coefficient, which signifies the sigmoidicity of the curve, and overall effect, indicated by dose concentration for a specified (e.g., 50%) response^14,15^. When there are extreme outcome observations, the improvement of robust beta regression in estimation accuracy could be substantial. Moreover, simulation studies further illustrate that the proposed approach provides a narrower confidence interval, which in turn suggests a higher efficiency to achieve better power in statistical testing even without acquiring additional experimental data. Illustrative examples using real-world data for cancer research and SARS-CoV-2 treatment are provided. The analyses are implemented using the freely accessible web-based application REAP, developed based on the Shiny package of R language, with which research scientists could conveniently upload their drug experiment dataset and perform the data analysis.

## Results

### REAP Shiny App

We developed a user-friendly analytic tool, coined “REAP” (Robust and Efficient Assessment of Potency), for convenient application of the robust dose-response estimation to real-world data analysis. It is established in an agile modeling framework under the parameterization of the beta law to describe a continuous response variable with values in a standard unit interval (0,1). We further exploited a robust estimation method of the beta regression, named the minimum density power divergence estimators (MDPDE)^13^, for dose-response estimation, with the tuning parameter optimized by a data-driven method^21^. The technical details are provided in the **Methods**.

REAP presents a straightforward analytic environment for robust estimation of dose-response curve and assessment of key statistics, including implementation of statistical comparisons and delivery of customized output for graphic presentation (**Figure 3**). The dose-response curve is a time-honored tool to convey the pharmacological activity of a compound. Through dose-response curves, we can compare the relative activity of a compound on different assays or the sensitivity of different compounds on an assay. REAP aims to make this job simple, estimation efficient, and results robust.

**Figure 3.**
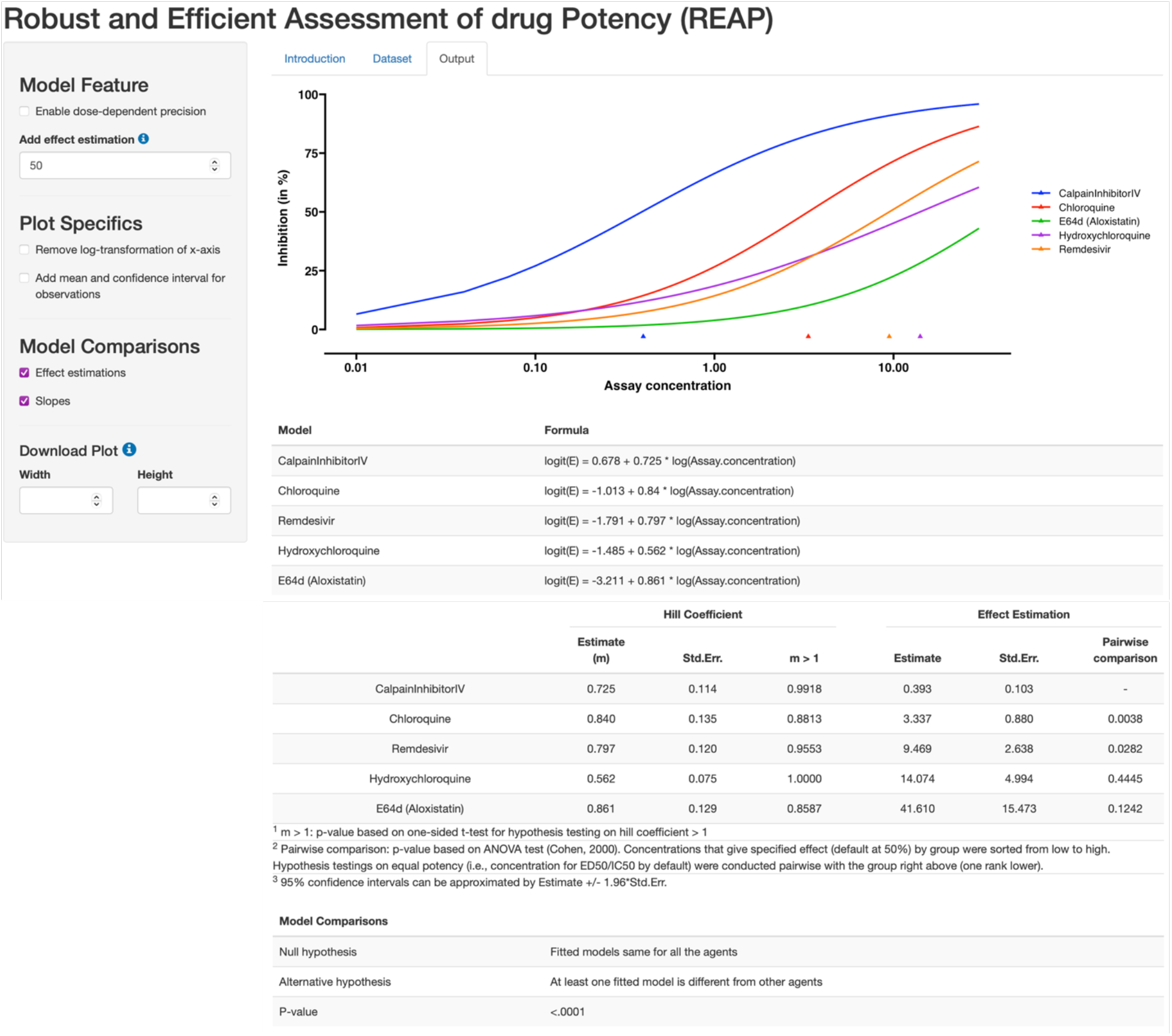
REAP App interface, with a highlight of Output section. Using the robust beta regression method, REAP produces a dose-response curve plot with effect and model estimations. The left panel allows users to specify model features and design plot specifics. REAP also provides hypothesis testing results to compare effect estimations, slopes and models.

There are three sections in REAP: Introduction, Dataset and Output. Users can have both overview and instruction of REAP in the Introduction. Dataset is uploaded in the Dataset section. The input dataset is mandated to be in a csv file format and contains three columns of data respectively for drug concentration, response effect and group name, in a specific order. It is recommended that users normalize the response variable to the range of (0,1) by themselves. Otherwise, REAP automatically will truncate the values exceeding the boundaries to (0,1) using a truncation algorithm (see **Supplementary Notes**). In the Output section, it generates a dose-response plot, along with tabulation for effect and model estimations. We also enable hypothesis testing for comparisons of effect estimations, slopes and models (i.e., comparing both intercepts and slopes) (see **Methods**). By default, the x-axis of the dose-response plot is log-scaled. In the plot, users can choose to add mean values and confidence intervals for data points under the same agent and dose level. Both plots and estimation tables are downloadable on REAP to plug in presentations and manuscripts for result dissemination.

The open-sourced REAP is freely available and accessible at https://xinying-fang.shinyapps.io/REAP/. We demonstrated it in two real-world examples, after presenting the simulation results, to illustrate the functionality of REAP.

### Simulations

We conducted simulation studies to investigate the robust beta regression model, in comparison to the linear regression model with data transformation, to characterize the median-effect equation under different scenarios. Details on the simulation setting are described in the **Supplementary Notes**.

**Figure 4** shows the results with data simulated using normal error terms. When the standard deviation (SD) is set to 0.005 under constant precision parameter setting, which refers to well-controlled experiments with only small systematic error, the point estimations of IC_50_, IC_90_, *β*_1_ and *β*_0_ are close to the pre-defined true values under all scenarios when using the robust beta regression. Compared to the linear regression, the robust beta regression shows small estimation bias and estimation error, and reasonable coverage probability in the estimates of IC_50_, IC_90_, *β*_1_ and *β*_0_ (**Supplementary Table 1)**. Meanwhile, the 95% CIs are much narrower (**Figure 4**), especially under the condition when data includes extreme values, indicating the efficiency of robust beta regression model in dose-response estimation. As SD increases, which hints the experiments may contain more errors, robust beta regression consistently performs well in estimating median-effect equation, considering the small bias and error in point estimation to true values and narrower 95% CI. For the scenarios where data do not include extreme values, robust beta regression is still better with improvement to a lesser extent. Lastly, substantive improvements are also observed when variances of error are non-constant but dose-dependent.

**Figure 4.**
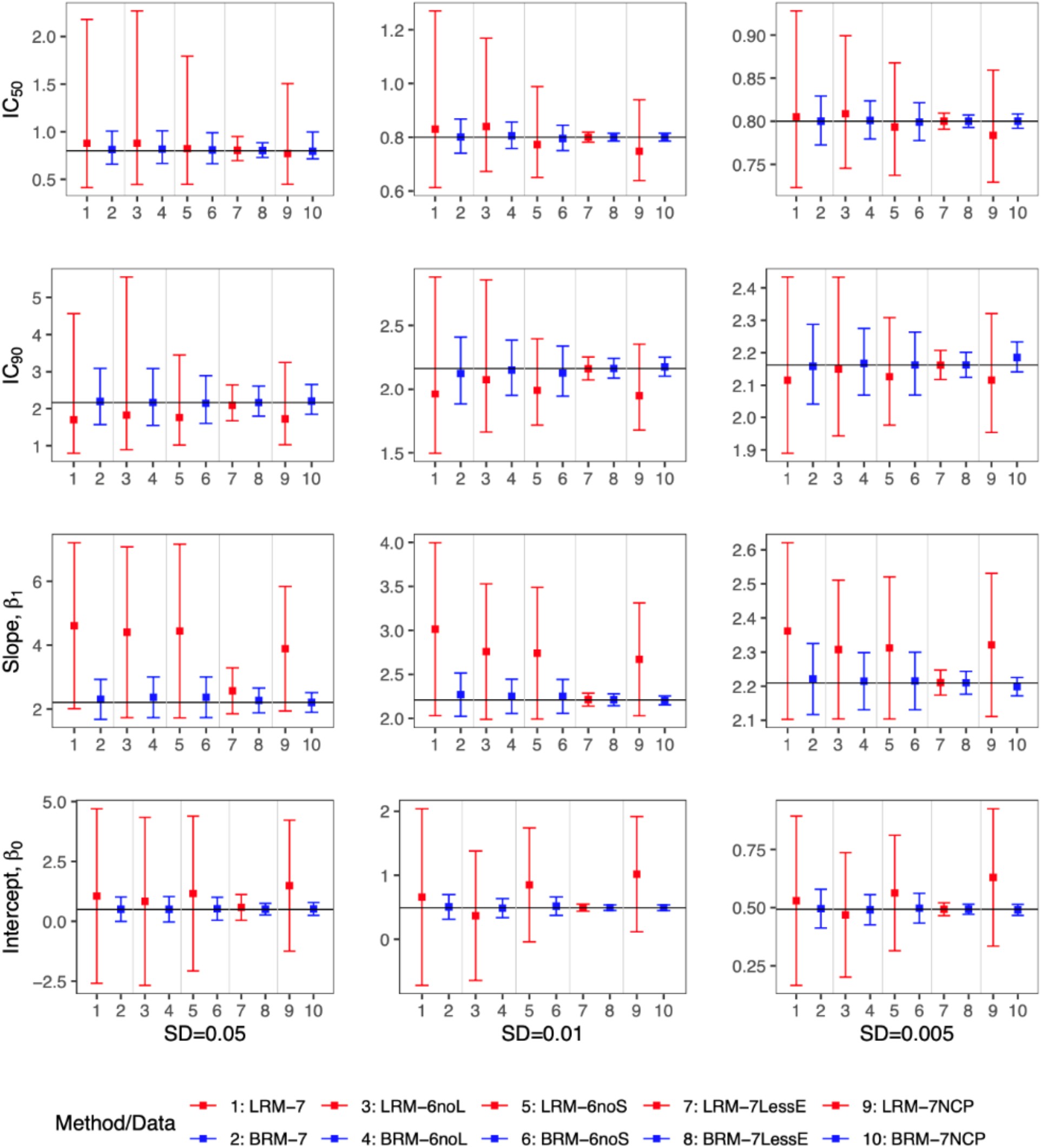
Comparison of the point estimates and 95% confidence intervals using linear regression model and robust beta regression model, with data simulated from normal error term. The vertical solid lines indicate the true values. The point estimation by robust beta regression was consistently closer to the true value with a narrower 95% CI compared to the linear regression model. LRM-7: LRM under 7 dose dataset with extreme data points; LRM-6noL: LRM under 6 dose dataset after removing the highest dose data point; LRM-6noS: LRM under 6 dose dataset after removing the lowest dose data point; LRM-7lessE: LRM under 7 dose dataset with less extreme data points; LRM-7NCP: LRM under 7 dose dataset with extreme data points and dose-dependent precision; BRM-7: BRM under 7 dose dataset with extreme data points; BRM-6noL: BRM under 6 dose dataset after removing the highest dose data point; BRM-6noS: BRM under 6 dose dataset after removing the lowest dose data point; BRM-7lessE: BRM under 7 dose dataset with less extreme data points; BRM-7NCP: BRM under 7 dose dataset with extreme data points and dose-dependent precision.

In parallel, similar results are obtained consistently with data simulated using beta error term, which induces heteroscedasticity (smaller on the two ends and bigger in the middle) at different dose levels (**Supplementary Figure 1, Supplementary Table 2**). All the results above demonstrate the sensitivity of standard regression model in dealing with datasets including extreme values. In addition, the result comparisons between the seven-dose set and the six-dose set with the largest or smallest dose eliminated display the potential worse influence of deleting extreme values directly in modeling dose-response using linear regression, which further notarizes the robustness and efficiency of the proposed robust beta regression.

Overall, the simulation study suggests that the robust beta regression model produces well-calibrated dose-response curves while being more robust and powerful than the standard regression model in estimating the median effect equation.

### B-cell lymphoma data

The first example of REAP application is dose-response curve estimation of the same agent under different cell lines. The data was originally from a study on using a drug called auranofin in treating B-cell lymphomas such as relapsed or refractory mantle cell lymphoma (MCL)^24^. As an FDA-approved for treatment of rheumatoid arthritis, auranofin targets thioredoxin reductase-1 (Txnrd1), and was repurposed as a potential anti-tumor drug to effectively induce DNA damage, reactive oxygen species (ROS) production, cell growth inhibition, and apoptosis in aggressive B-cell lymphomas, especially in TP53-mutated or PTEN-deleted lymphomas.

In the experiment, the effect of auranofin was evaluated in six MCL cell lines (Z-138, JVM-2, Mino, Maver-1, Jeko-1, and Jeko-R) with auranofin in concentrations ranging from 0 to 5 μM for 72 h and tested cell viability using a luminescent assay. The confidence interval bars of observed dose-response in **Figure 5** show that the sample variance of error from repeated measurements decreased with the increase of auranofin concentrations. To account for the heteroscedasticity and asymmetry in the variance, we enable a dose-dependent precision (proportional to inverse variance) in REAP, adding log(*dose*) as an additional regressor for the precision parameter. **Figure 5** shows the fitted dose-response curves with the dose-dependent precision. The test for homogeneity (p-value < 0.0001) suggests distinct dose-response between cell lines. The estimation of intercepts, hill coefficients and pairwise comparisons of IC_50_ estimations are provided in **Supplementary Table 3**.

**Figure 5.**
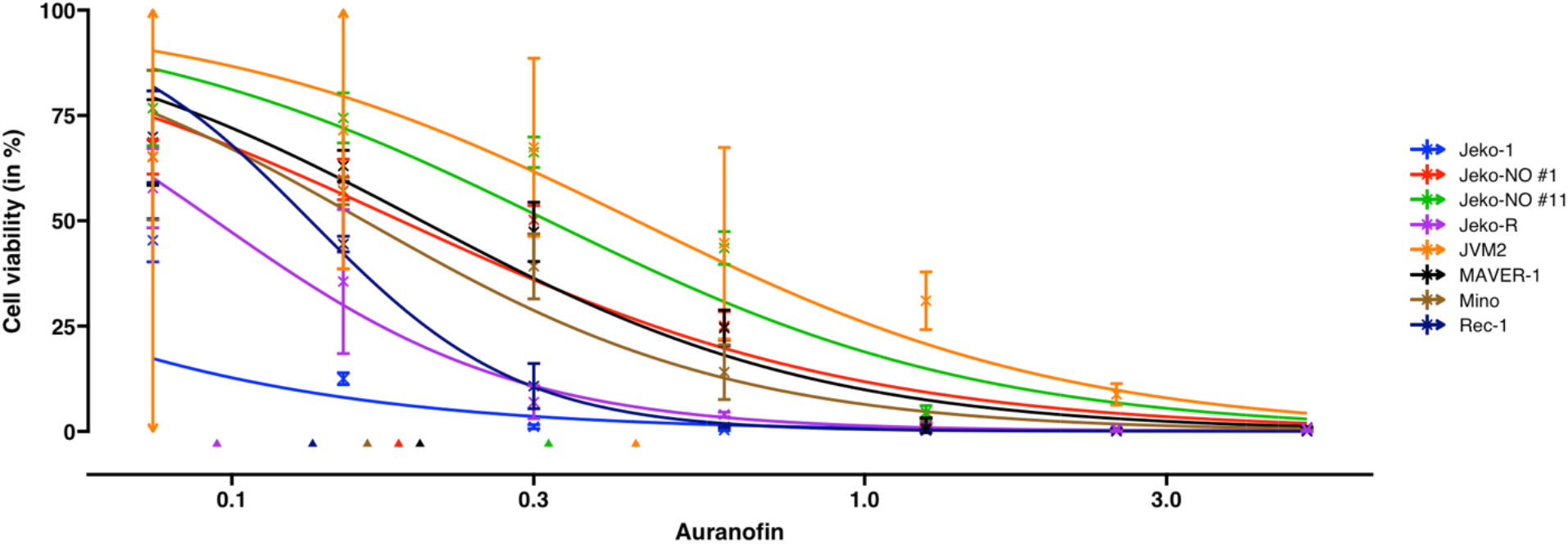
Dose-response curve estimation of auranofin (μM) under different MCL cell lines. The doseresponse curve was fitted with a dose-dependent precision with log (*dose*) as an additional regressor for the precision estimator. Observed dose effects are displayed with 95% confidence interval bars, which end with arrows when estimated confidence intervals exceed (0,1). Triangles at the bottom indicate IC_50_ values for each MCL cell line. MCL: mantle cell lymphoma.

### SARS-CoV-2 data

The second example is on the dose-response curve estimation in antiviral drug development for coronavirus disease 2019 (COVID-19). At the beginning of 2020, COVID-19 broke out at an unprecedented pace internationally, but there were limited therapeutic options for treating this disease. Therefore, many compounds and their combinations were rapidly tested *in vitro* against the SARS-CoV-2 virus to identify potentially effective treatments and prioritize clinical investigation.

In the data^25^, the benchmark compound collection consists of five known antivirals, including remdesivir, E64d (aloxistatin), chloroquine, calpain Inhibitor IV and hydroxychloroquine. The *in vitro* experiment was performed using the same biological batch of SARS-CoV-2 virus and conducted in biosafety level-3. In the original publication^25^, the dose-response curves were fitted by linear regression, which could yield inconclusive estimation (e.g., hydroxychloroquine in Fig. 1G of Bobrowski et al. (2020)^25^), while the estimated inhibition tends to exceed 1 when concentration is larger than 10 *μ*M. REAP gives reasonable estimation for the dose-response curves (**Figure 6**). The hypothesis testing results show that at least one slope estimation is different from other antivirals (p-value = 0.0003) and at least one EC_50_ estimation is different from others (p-value = 0.003). Calpain Inhibitor IV shows a higher potency than other agents including hydroxychloroquine (p-value = 0.0038, **Supplementary Table 4**).

**Figure 6.**
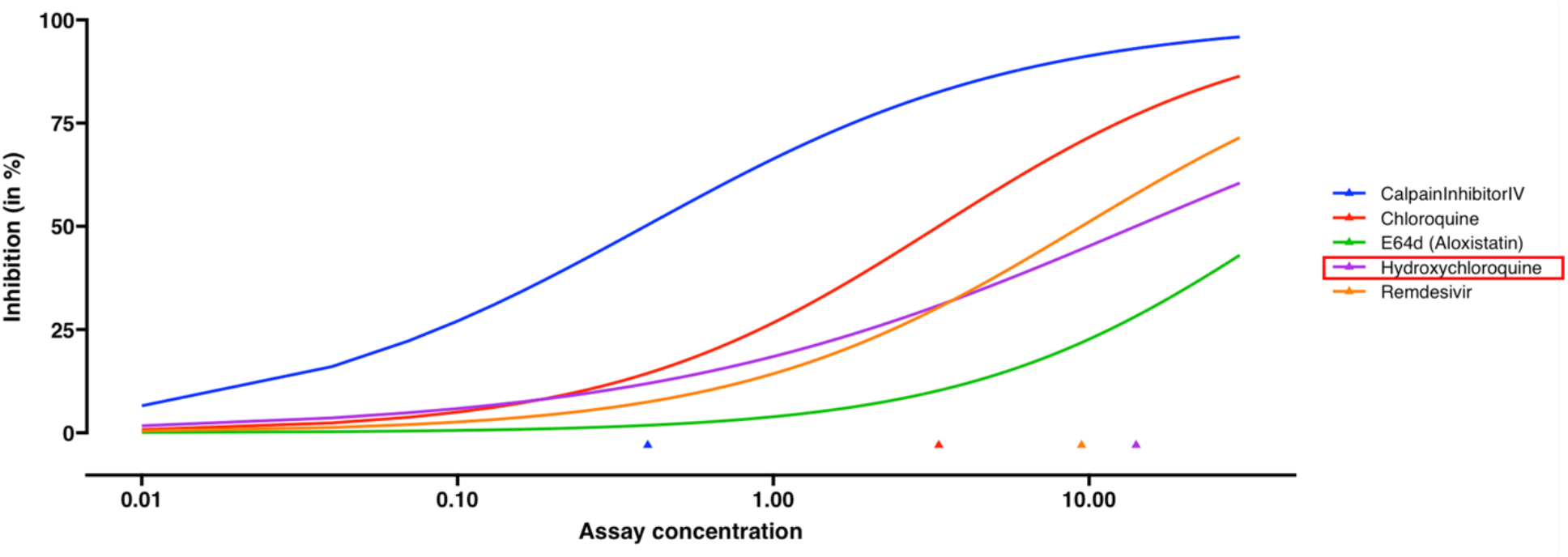
Dose-response curve estimation of anti-viral drugs under the same biological batch with SARS-CoV-2 data. The robust beta regression gives reasonable estimations to dose-response curve of hydroxychloroquine, compared to the inconclusive dose-response curve fitted by linear regression in Bobrowski et al. (2020). The plot is generated without selecting the option of mean and confidence interval for observations. Triangles indicate the estimated EC_50_ values for each drug.

## Discussion

Quantifying the potency of a compelling substance is always a central topic in life sciences^26^. It is a vital component of research in pharmacology, but also prevalent in the fields of toxicology, environmental science, agrochemistry, and medicine, among many others. For instance, the description of dose-response curves can provide the initial toxicological risk assessment^27^, and guide *in silico* modeling of toxic doses to humans and the environment^28^. Based on proper identification of dose-response relationship from *in vitro* assays, studies can successfully predict systemic toxicological effects *in vivo* without additional *in silico* modelling^29^. Nevertheless, it necessitates accurate and reliable description of the dose-response curve, which further demands robust and efficient modeling strategies to account for embedded variability in observed response and to derive solid inference with valid quantification of uncertainty.

The dose-response estimation could be substantially biased by the standard regression modeling. In the illustrative example (**Figure 1**), the estimated IC_50_ dose indeed effects the 70% fraction of cell affected, while the estimated response at the true IC_50_ dose is only 22%. Such a large discrepancy is sourced by a small (<0.5%) single measurement error, which is common and inevitable in any regular *in vivo* experiment, but could engender profound impact to assessment of drug potency and determination of synergy in drug combinations. In addition, the modeling strategy of deleting those extreme values (e.g., **Figure 2**, or 6noL and 6noS datasets in **Figure 4 and Supplementary Figure 1**) is futile to improve the poor performance of standard regression model, but may further impair the estimation efficiency and accuracy. In general, it fails to reduce bias but only introduces larger uncertainty in estimation of dose concentration, especially at extreme responses (e.g., IC_90_).

We develop REAP for assessment of drug potency to address concerns in this regard. It has substantial advantages over existing methods by reducing the impact of random errors due to implicit variations in the experimental data. To our best knowledge, it is also for the first time that beta regression is introduced to dose-response estimation. The underlying modified robust beta regression model estimated by the data-driven tuning parameter is resilient to estimation bias caused by extreme observations, which is a routinely encountered situation for deficient dose-response estimation using the standard estimation approach. The proposed approach is also efficient in quantitative characterization of dose-response curves with narrower confidence intervals for key estimators. Furthermore, REAP can simultaneously model the data heterogeneity with a dose-dependent precision component (**Figure 5**). It is simply different from other dose-response methods, in which a vector of weights have to be (possibly mis-)specified externally. REAP is an open-source and user-friendly platform, developed for diverse non-computational scientists for hands-on wet-laboratory data analysis in regular use, and can be hosted within R shiny environment under Windows, Linux, and Mac system or deployed in Docker available as a web server.

Our work potentially can be useful in applications of drug screening. The proposed method and the developed REAP App allow for the robust and efficient estimation and accounting for outliers as well, making it fitted particularly in a high-throughput setting. As the result of a complex and dynamic cascade of events, exposure time is another important factor ultimately affecting the dose-response. For *in vitro* experiments measured at different time-points in a choice of cell-lines and expressed by a variety of assays^30^, the proposed modeling framework can be naturally extended to model time-dependent cytotoxicity while controlling for fixed or random effects. Furthermore, the application of robust and efficient dose-response estimation can be integrated into methods to identify drug interaction effect^6,31^. There is a venerable history that multi-agent combination therapies demonstrate great advantages in improving therapeutic efficacy and revolutionize patient outcomes in a wide range of diseases. Robust and efficient estimation of the dose-response curve would be crucial in investigation of adequate drug combinations.

The developed method has limitations. We presented a model of the median effect equation for doseresponse curve estimation based on mass action law. While in specific scenarios other laws may be considered more suitable to describe the biomedical systems, the current modeling framework can be naturally adapted for other dose-response functions like probit (via cumulative normal distribution) and Weibull model^32^, or any other continuous distribution functions. In addition, the median-effect equation to characterize pharmacological activity assumes the compound can affect all the cells. From a quantitative perspective, a compound that cannot reach high binding affinity will yield an overconservative estimation for median effective dose of a drug. However, in comparison to the sensitivity of different compounds in an assay, it is not harmful because the less effective compounds will be more easily identified. If it is a concern that the maximal effects of candidate compounds are different and the aim is to accurately model the dose-response curve, the Emax model could be a better choice^33^. Furthermore, the robust beta regression approach in REAP cannot handle values equal or less than 0, or equal or greater than 1. Thus, we developed a sequential data truncation algorithm in REAP to overcome the limitation of the conventional transformation *(y * (n−1) + 0.5) / n*, which could be too rough in doseresponse curve estimation particularly when the sample size *n* for each group is relatively small. Although empirically we have validated it using simulated data, the algorithm could be improved by future work to retain information more efficiently.

In summary, a good modeling strategy must effectively characterize the nature of the observed doseresponse pattern^16^. Rapid advances in novel drug development and considerable deficiency in modeling data with extreme values offer an appealing opportunity for next-generation quantitative approaches. While many aspects of the techniques discussed here fit in the statistical framework of robust beta regression, our aim is to clearly apply and rigorously customize the analytic considerations, to reduce bias and ameliorate efficiency in routinely used dose-effect estimation, and to facilitate the convenient analytic implementation and dissemination. Experimental conditions and candidate drug potency could inevitably vary in practice, but REAP provides a great tolerance for points with extreme values, solid support for accurate and efficient dose-response curve estimation, and useful reference to the future development of methodology in drug investigation. Overall, we anticipate that our work will contribute more to quantitative analysis in assessment of drug potency in pre-clinical research.

## Methods

### Median-effect equation and dose-response curve

The median-effect equation describes a popular model of the dose-response relationship based on the median effect principle of the mass action law in various biological systems^2^. Assume *f_a_* and *f_u_* are the fractions of the system affected and unaffected by a drug concentration *d*. The median-effect equation states that

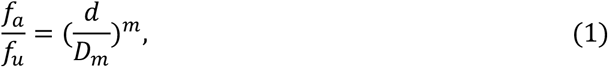

where *m* is the Hill coefficient signifying the sigmoidicity of the dose-effect curve and *D_m_* is the dose of a drug required to produce the median effect, which is analogous to the more familiar *IC*_50_ (drug concentration that causes 50% of the maximum inhibitory effect), *ED*_50_ (half-maximum effective dose), or *LD*_50_ (median lethal dose) values ^13^. For example, if an inhibitory substance is of interest, the parameter *m* measures the cooperativity in the binding of multiple ligands to linked binding sites, and the parameter *D_m_* = *IC*_50_, defined by the concentration that causes 50% of the maximum inhibitory effect.

Given *f_a_* + *f_u_* = 1, the median-effect equation (1) is equivalent to

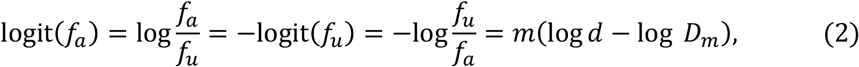

where logit(*p*) denotes the logit function 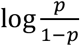. The equation (2) shows a log-linear relationship between the drug dose *d* and its effect *f_a_* (or *f_u_*, if it is, for example, the % survival of interest) after a logit transformation. Because from a modeling perspective the identical strategy can be applied to model both *f_a_* and *f_u_*, for the effect on cell fraction *E*, we can re-write equation (2) to be:

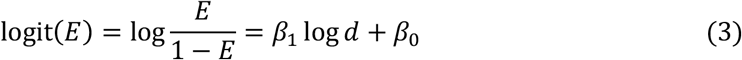

where *β*_0_ is the intercept and *β*_1_ the slope of the response curve. In this presentation, the median effect dose

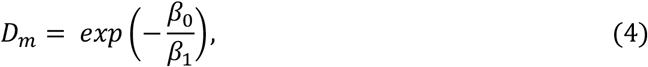

the Hill coefficient

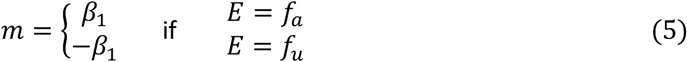

and the dose-response curve

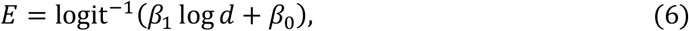

where 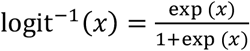 is the inverse-logit function.

### Beta regression model for dose-response curve estimation

We will review the beta regression model which for the first time will be applied in dose-response estimation. The effect *E* and the parameters *β* = (*β*_0_, *β*_1_) in equation (3) cannot be directly observed, but they can be estimated using experimental data, in which the observed sample cell fraction *y* produced by the drug dose *d* is a random variable with mean *E*. It is clear that effective estimation must properly account for random variation and be based upon a model that not only matches the nature of the response variable, but adequately characterizes the observed dose-response pattern^16^.

Among all the unknown quantities, the parameters *β* could be first estimated and play a fundamental role in supporting the inference for others. In the standard estimation procedure based on linear regression, 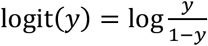 is regressed on log *d* to get the inference on parameters *β*. Subsequently, the doseresponse curve can be estimated by equation (6), and (*D_m_, m*) can be derived based on equations (4) and (5) for median-effect equation (2). Because the extreme values of *y* close to 0 or 1 could yield very large values of logit(*y*) (approaching to −∞ or +∞, respectively, if *y* → 0 or 1), and induce significant bias in estimation of *β*, the accuracy of the estimated dose-response curve and median-effect equation is in question when there exist extreme values in the dataset.

The beta regression model describes a response variable *y* with continuous values restricted to the open standard unit interval^17,18^. In a classic beta regression framework, the beta regression model uses a parameterization of the beta law that is indexed by the mean parameter *μ*, and the precision parameter *ϕ* that controls the overall variation^12^. To model the dose-response relationship for the cell fraction *E*, we assume that the response *y* is a beta-distributed random variable and its mean *μ* = *E* has the form of equation (6), where *d* is the dose producing effect *E, β*_1_ and *β*_0_ are the regression parameters. Estimation of regression parameters *β* can be performed using maximum likelihood method to derive point estimate 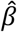 and covariance matrix *Σ*.

Beta regression is resistant to extreme values and provides reliable estimations (**Fig. 1**). Compared with the standard approach, which applies a non-linear transformation in the response for an approximation to the normal distribution, the beta density can take on a variety of shapes to account for non-normality and skewness^19^. In the presence of heteroskedasticity and asymmetry, two common problems frequently observed in limited range continuous response data, an empirical study showed that the beta regression provided the best estimation among several alternatives^20^.

### Robust beta regression model with MDPDE

We will present a modified robust beta regression approach in REAP implementation, which is established based on density power divergence for robust estimation^13^, but further improved after we introduce a data-driven method to identify the optimal tuning parameter. The standard beta regression potentially could still be sensitive against outliers because its inference is based on the maximum likelihood estimation. Ghosh^13^ developed the robust minimum density power divergence estimators (MDPDE) that address the problem by minimizing the average density power divergence (DPD)

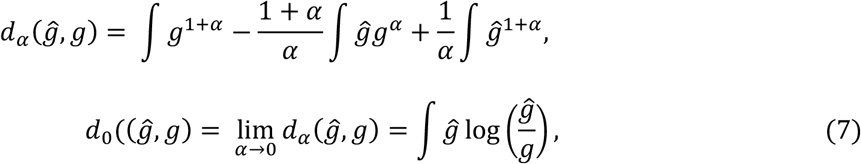

between the empirical density 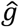 and the beta model density function *g* ≡ *Beta*(*μϕ*, (1 − *μ*)*ϕ*). *α* is a non-negative tuning parameter, smoothly connecting the likelihood disparity (at *α* = 0) to the *L_2_*-Divergence (at *α* = 1). The parameter of interest *β* is estimated by minimizing the DPD measure between *g_i_* and the density, 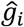, estimated by data:

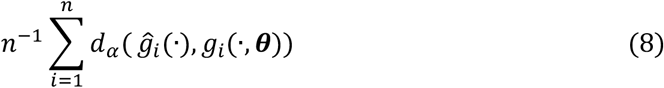

or equivalently, minimizing the objective function using the estimation equations:

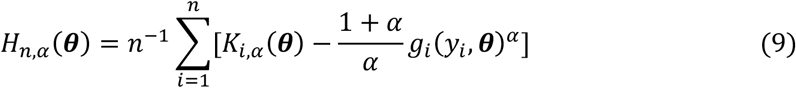

where 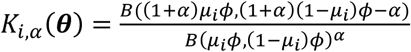.

MDPDE improves the standard beta regression with the DPD measure and a fixed tuning parameter. The recommended *α* is around 0.3 to 0.4, but simply assigning a fixed *α* in [0.3,0.4] is not applicable in many cases. Here we adopted a data-driven method^21^ to identify the optimal *α*. The search for the optimal *α* starts with a grid of *α* and a pre-defined *α_max_* and grid size *p*, which generates a sequence of equally spaced 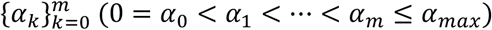. MDPDE calculates the corresponding ***θ*** and *se*(***θ***) with each *α* so that we get a vector of standardized estimates:

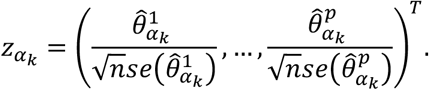

The standardized quadratic variations (SQV) are defined by:

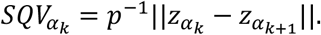

We compare each *SQV_α_k__* with a pre-defined threshold *L* (*L* > 0). If all *α_k_* satisfy the stability condition of *SQV_α_k__* < *L*, then the optimal *α* equals the minimal *α* in *α_k_*. Otherwise, restart the search with a new grid of *α_k_*. The new grid of the same size *p* is picked from the sequence 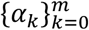 starting from the largest *α_k_* that fails the stability condition. Repeat searching until the current grid satisfies the stability condition or *α_max_* is reached. If the stability condition is satisfied before *α_max_* is reached, then optimal *α* equals the minimal value in the grid of *α_k_*. If *α_max_* is reached, then optimal *α* equals 0, which is equivalent to the maximum likelihood estimation.

### Point estimate and its confidence interval for drug activity measurements

The objective of analysis is to characterize the dose-response curves in equation (2) and quantify in vitro drug potency. Popular drug activity measurements include Hill coefficient *m* and median effect dose *D_m_*. In some circumstances, other measurements such as instantaneous inhibitory potential (*IIP*), which directly quantifies the log decrease in single-round infection events caused by a drug at a clinically relevant concentration, are of special interest^22^.

The MDPDE for beta regression model provides a robust strategy to estimate *β*, from which the point estimates and confidence intervals of relevant drug activity measurements can be derived. Mathematically, those drug activity quantities can be written as functions of parameters *β* with an explicit form. Subsequently, their point estimates and confidence intervals can be derived based on the inference of *β*. For example, given a point estimate 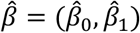, the point estimate for 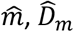 as a single value, and 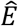 as a function of dose *d* can be computed using equations (4 – 6).

It is important to construct the confidence interval around the point estimate to gauge the estimation uncertainty. With different levels of measurement error from either well-managed or lousy experiments, the levels of evidence vary for statistical inference, even if it is derived the same point estimates for the intercept *β*_0_, slope *β*_1_ and the corresponding dose-response curve. Given the point estimate 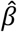 and its positive-definite covariance matrix *Σ* to account for variability in observed response, we apply the multivariate delta method and approximate the variance estimate after assuming asymptotic normality^23^. As demonstrated in our simulation studies, the constructed (1 − *α*) × 100% confidence interval consistently provides better results to quantify the (1 − *α*) × 100% coverage probability. More importantly, the width of the constructed confidence interval was narrower than that from a linear regression model, suggesting that our approach is more efficient with a higher statistical power (**Supplementary Tables 1-2**).

### Comparison of the dose-response curves

When we estimate multiple dose-response curves with the data collection experiments conducted in a similar setting, it is often of interest to statistically compare the drug potency and/or Hill coefficients. A typical comparison may occur when we examine the similarity of response from different drugs, explore the additional effect of a drug combined with certain monotherapy, or assess the homogeneity of a drug to different patient samples or cell lines. In the beta regression framework, the statistical comparison can be conducted by first comparing independent fits for each curve with a global fit that shares the common parameters among different groups. Subsequently, the likelihood ratio test can be applied to examine whether the same Hill coefficient or one dose-response curve can adequately fit all the data. The only exception is to assess whether median effect doses are the same in different groups, while an F test is used for the single parameter testing. If the global test for potency shows a significant p-value, a pairwise comparison can be conducted using two-sided t-test for the ordered groups with Benjamini-Hochberg correction for multiplicity.

## References

1. Chou, T. C. Theoretical basis, experimental design, and computerized simulation of synergism and antagonism in drug combination studies. Pharmacological Reviews vol. 58 621–681 (2006).

2. Chou, T. C. Derivation and properties of Michaelis-Menten type and Hill type equations for reference ligands. J. Theor. Biol. 59, (1976).

3. Chou, T. C. & Talalay, P. Quantitative analysis of dose-effect relationships: the combined effects of multiple drugs or enzyme inhibitors. Adv. Enzyme Regul. 22, 27–55 (1984).

4. Chou, T. & Rideout, D. C. The median-effect principle and the combination index for quantitation of synergism and antagonism. in Synergism and Antagonism in Chemotherapy (1991).

5. Greco, W. R., Bravo, G. & Parsons, J. C. The search for synergy: a critical review from a response surface perspective. Pharmacol. Rev. 47, (1995).

6. Lee, J. J. & Kong, M. Confidence Intervals of Interaction Index for Assessing Multiple Drug Interaction. Stat. Biopharm. Res. (2009) doi:10.1198/sbr.2009.0001.

7. Roell, K. R., Reif, D. M. & Motsinger-Reif, A. A. An introduction to terminology and methodology of chemical synergy-perspectives from across disciplines. Front. Pharmacol. 8, 1–11 (2017).

8. Gadagkar, S. R. & Call, G. B. Computational tools for fitting the Hill equation to dose-response curves. J. Pharmacol. Toxicol. Methods 71, 68–76 (2015).

9. Solzin, J. et al. Action limit outlier test: A novel approach for the identification of outliers in bioassay dose-response curves. Bioanalysis 12, 1459–1468 (2020).

10. Schmidheiny, K. Heteroskedasticity in the Linear Model. Econometrica (2009).

11. Williams, J. D., Birch, J. B., Woodall, W. H. & Ferry, N. M. Statistical monitoring of heteroscedastic dose - Response profiles from high-throughput screening. J. Agric. Biol. Environ. Stat. 12, 216–235 (2007).

12. Ferrari, S. L. P. & Cribari-Neto, F. Beta regression for modelling rates and proportions. J. Appl. Stat. 31, 799–815 (2004).

13. Ghosh, A. Robust inference under the beta regression model with application to health care studies. Stat. Methods Med. Res. 28, 871–888 (2019).

14. Shen, L. et al. Dose-response curve slope sets class-specific limits on inhibitory potential of anti-HIV drugs. Nat. Med. 14, 762–766 (2008).

15. Sampah, M. E. S., Shen, L., Jilek, B. L. & Siliciano, R. F. Dose-response curve slope is a missing dimension in the analysis of HIV-1 drug resistance. Proc. Natl. Acad. Sci. U. S. A. 108, 7613–7618 (2011).

16. Lyles, R. H., Poindexter, C., Evans, A., Brown, M. & Cooper, C. R. Nonlinear model-based estimates of IC50 for studies involving continuous therapeutic dose-response data. Contemp. Clin. Trials 29, 878–886 (2008).

17. Johnson, N., Kotz, S. & Balakrishnan, N. Continuous univariate distributions, volume 2. (1995).

18. Simas, A. B., Barreto-Souza, W. & Rocha, A. V. Improved estimators for a general class of beta regression models. Elsevier https://www.sciencedirect.com/science/article/pii/S0167947309003107 (2008).

19. Smithson, M. & Verkuilen, J. A better lemon squeezer? Maximum-likelihood regression with beta-distributed dependent variables. Psychol. Methods (2006) doi:10.1037/1082-989X.11.1.54.

20. Kieschnick, R. & Mccullough, B. D. Regression analysis of variates observed on (0, 1): Percentages, proportions and fractions. Stat. Model. 3, 193–213 (2003).

21. Ribeiro, T. K. A. & Ferrari, S. L. P. Robust estimation in beta regression via maximum Lq-likelihood. (2020).

22. Shen, L., Rabi, S. A. & Siliciano, R. F. A novel method for determining the inhibitory potential of anti-HIV drugs. Trends Pharmacol. Sci. 30, 610–616 (2009).

23. Bickel, P. & Doksum, K. Mathematical statistics: basic ideas and selected topics, volumes I-II package. (2015).

24. Wang, J. et al. Repurposing auranofin to treat TP53-mutated or PTEN-deleted refractory B-cell lymphoma. Blood Cancer J. 9, (2019).

25. Bobrowski, T. et al. Synergistic and Antagonistic Drug Combinations against SARS-CoV-2. Mol. Ther. 29, 873–885 (2021).

26. Schindler, M. Theory of synergistic effects: Hill-type response surfaces as ‘null-interaction’ models for mixtures. Theor. Biol. Med. Model. 14, 1–16 (2017).

27. Council, N. R. Toxicity testing in the 21st century: A vision and a strategy. Toxicity Testing in the 21st Century: A Vision and a Strategy (National Academies Press, 2007). doi:10.17226/11970.

28. Blaauboer, B. J. et al. The use of biomarkers of toxicity for integrating in vitro hazard estimates into risk assessment for humans. in Altex vol. 29 411–425 (Elsevier GmbH, 2012).

29. Groothuis, F. A. et al. Dose metric considerations in in vitro assays to improve quantitative in vitro-in vivo dose extrapolations. Toxicology 332, 30–40 (2015).

30. Byrne, H. J. & Maher, M. A. Numerically modelling time and dose dependent cytotoxicity. Comput. Toxicol. 12, 100090 (2019).

31. Lee, J. J., Kong, M., Ayers, G. D. & Lotan, R. Interaction index and different methods for determining drug interaction in combination therapy. J. Biopharm. Stat. 17, 461–480 (2007).

32. Christensen, E. R. Dose-response functions in aquatic toxicity testing and the Weibull model. Water Res vol. 18 (1984).

33. Jack Lee, J., Lin, H. Y., Liu, D. D. & Kong, M. Emax model and interaction index for assessing drug interaction in combination studies. Front. Biosci. - Elit. 2 E, 582–601 (2010).

