## Supplementary materials for "Robust and Efficient Assessment of Potency (REAP): A Quantitative Tool for Dose-response Curve Estimation"

### Supplementary Information

Content

Truncation Strategy…………………………………………………………………………..…2

Data Simulating Process.………………………………………………………………..…3

Supplementary Figure 1…………………………………………………………………..…5

Supplemental Table 1……………….…………………………………………………………..7

Supplemental Table 2…………….………………………………………………………..…8

Supplemental Table 3…….……..…………….………………………………………..…9

Supplemental Table 4……….………………………………………………………...10

**Truncation strategy**

Based on the median-effect equation method by Chou TC, the software “CompuSyn” was published. In the data entry illustration of this software, they pointed out the sensitivity limits of data points, e.g., too low (fa<0.02) and too high (fa>0.99) and suggested that such data points out of effect may be edited or deleted.

There are some data truncation algorithms in the literature. Two obvious remedies are proportionally “shrinking” the range to a sub-range nearly covering the unit interval (e.g., [.00001, .99999]) or simply adding a small amount to 0-valued observations and subtracting the same amount from 1-valued observations while leaving the other observations unchanged. MacMillan and Creelman (2005) mentioned a method which is frequently used in practice in areas such as signal detection is to add 1/2n to a 0 observation and subtract 1/2n from a 1 observation, where n is the total number of observations. Besides, Smithson M, Verkuilen J (2006) demonstrated that a useful transformation in practice is (y * (n−1) + 0.5) / n, which is also mentioned by the documentation for R Betareg package for conditions when data assumes the extremes 0s and 1s. In dose-response curve estimation, this treatment could be too rough, especially when n is small.

To minimize the impact from truncation of data points, we apply the following algorithm. The first step is to shrink the data range to [1e-9, 1-1e-9]. If there still exist abnormal conditions, we will sequentially shrink the data range of abnormal ones to [1e-8, 1-1e-8], …, until [1e-3, 1-1e-3] or noon-exist of abnormal conditions. Then, if it still exists, though rarely, the transformation of (y * (n−1) + 0.5) / n, where n is the sample size, in the documentation of R Betareg package would be applied. We have conducted simulations to test this algorithm in various scenarios with difference errors and it achieved reasonable performance in handling all conditions.

**Data simulating process**

In the simulation study, both robust beta regression and linear regression are applied to estimate dose-response curves under different scenarios. The point estimations and 95% confidence intervals of $IC_{50}$, $IC_{90}$, $\beta_0$ and $\beta_1$ under each method will be obtained and then, be compared to evaluate the model performance.

To generate data for simulation studies, we define the dose set for simulation as 0.1, 0.2, 0.4, 0.8, 1.6, 3.2 and 6.4 $\mu M$, which consists of 7 doses, and choose the appropriate true curve with $\beta_1 = 2.2098$ and $\beta_0 = 0.4931$ such that the corresponding effects of the smallest and largest dose are 0.01 and 0.99, respectively. Let's call the true curve "$E = f(\log(dose))$". Then, the following equation is applied to generate data by inducing random error into effect:

$$E = true + error = f(\log(dose)) + error$$

We simulated data with two types of errors, normal error term and beta error term, to examine the accuracy and sensitivity of model performance in general setting. The normal error term is implemented with different standard deviations (SDs), e.g. 0.005, 0.01 and 0.05, while the beta error term is under different precision parameter $\phi$, e.g. 35, 15, 5. Note that the larger the $\phi$, the smaller the variance. By implementing under different SD or $\phi$, it allows for generation of not only well-controlled data which is assumed for experiments with almost no error, but also noised data which is more identical to real world data. The generated data is 1 replicate given each dose level with the total simulation size equal 10,000 for each choice of SD or $\phi$. Since the defined dose set is symmetric, we set up several scenarios under both error terms above: 1) full 7-dose set with extreme values; 2) 6-dose set after removing the largest dose; 3) 6-dose set after removing the smallest dose; 4) full 7-dose set with less extreme values by obtaining the smallest and largest dose levels with corresponding effect as 0.1 and 0.9 under the same true curve. The scenarios 1-4 assume constant precision parameter during data simulation and modeling process.

To mimic the real-world environment of data collections, the assumption of equal variance doesn't always hold. Thus, we also set up the $5^{th}$ scenario which uses full 7-dose set with extreme values with non-constant SD or precision parameter during data simulation and modeling process, but linearly dose-dependent. For normal error term, the modified SDs for data simulation have the form of $SD^* = (\gamma_0 + \gamma_1 * log.dose) * SD$; for beta error term, the modified precisions $\phi$ for data simulation have the form of $\phi^* = (\gamma_0 + \gamma_1 * log.dose) * \phi$. Assuming the same true dose-response curve as the previous simulation, we pre-defined $\gamma_0$ and $\gamma_1$ as 0.25 and 0.1378 such that the average of $SD^*$ is close to $SD$, and the average of $\phi^*$ is close to $\phi$, respectively.

**Supplementary Figure 1.** Comparison of the point estimates and 95% confidence intervals using linear regression model and robust beta regression model, with data simulated from beta error term. The vertical solid lines indicate the true values. The point estimation by robust beta regression was consistently closer to the true value with a narrower 95% CI compared to the linear regression model. LRM-7: LRM under 7 dose dataset with extreme data points; LRM-6noL: LRM under 6 dose dataset after removing the highest dose data point; LRM-6noS: LRM under 6 dose dataset after removing the lowest dose data point; LRM-7lessE: LRM under 7 dose dataset with less extreme data points; LRM-7NCP: LRM under 7 dose dataset with extreme data points and dose-dependent precision; BRM-7: BRM under 7 dose dataset with extreme data points; BRM-6noL: BRM under 6 dose dataset after removing the highest dose data point; BRM-6noS: BRM under 6 dose dataset after removing the lowest dose data point; BRM-7lessE: BRM under 7 dose dataset with less extreme data points; BRM-7NCP: BRM under 7 dose dataset with extreme data points and dose-dependent precision.

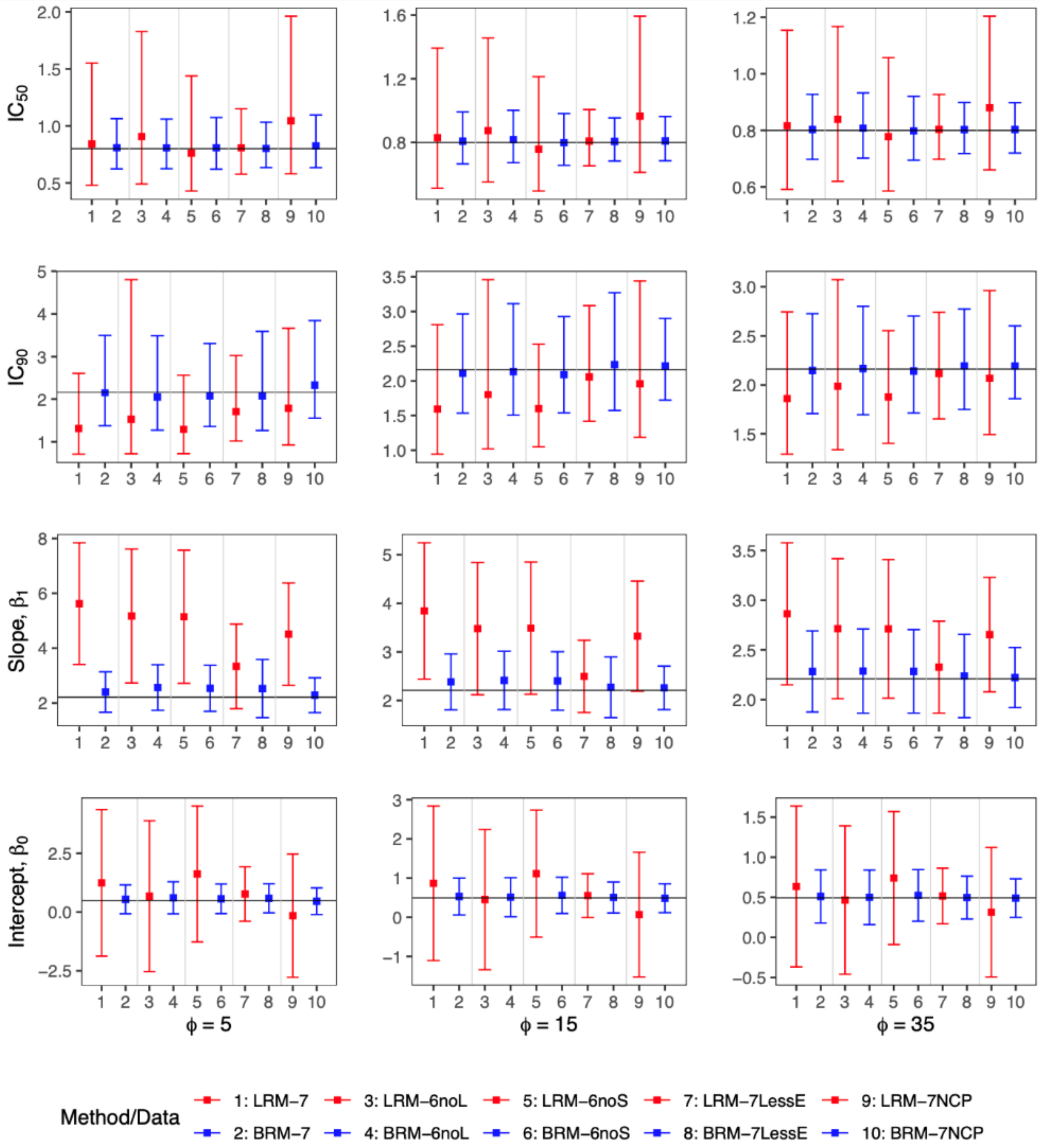

$IC_{50}$
$IC_{90}$
Slope, $\beta_1$
Intercept, $\beta_0$
$\phi = 5$
$\phi = 15$
$\phi = 35$
Method/Data
1: LRM−7
3: LRM−6noL
5: LRM−6noS
7: LRM−7LessE
9: LRM−7NCP
2: BRM−7
4: BRM−6noL
6: BRM−6noS
8: BRM−7LessE
10: BRM−7NCP

**Supplemental Table 1.** Simulation result of bias, RMSE and 95% CI coverage probability corresponding to normal error term constant precision

| Scenario | Method | Bias | | | | RMSE | | | | 95% CI Coverage Probability | | | |
|---|---|---|---|---|---|---|---|---|---|---|---|---|---|
| | | $IC_{50}$ | $IC_{90}$ | $\beta_0$ | $\beta_1$ | $IC_{50}$ | $IC_{90}$ | $\beta_0$ | $\beta_1$ | $IC_{50}$ | $IC_{90}$ | $\beta_0$ | $\beta_1$ |
| **(a) data simulated using normal error term with SD=0.005** | | | | | | | | | | | | | |
| 7 doses with extreme values | LRM | 0.005 | -0.047 | 0.037 | 0.152 | 0.098 | 0.298 | 0.525 | 0.557 | 0.954 | 0.773 | 0.943 | 0.666 |
| | BRM | 0.000 | -0.004 | 0.003 | 0.011 | 0.013 | 0.130 | 0.044 | 0.088 | 0.981 | 0.927 | 0.975 | 0.889 |
| 6 doses after removing largest | LRM | 0.009 | -0.01 | -0.024 | 0.098 | 0.045 | 0.184 | 0.129 | 0.517 | 0.967 | 0.921 | 0.971 | 0.691 |
| | BRM | 0.001 | 0.005 | -0.00 | 0.005 | 0.009 | 0.132 | 0.024 | 0.089 | 0.955 | 0.892 | 0.950 | 0.844 |
| 6 doses after removing smallest | LRM | -0.007 | -0.036 | 0.070 | 0.102 | 0.037 | 0.306 | 0.362 | 0.533 | 0.969 | 0.623 | 0.933 | 0.695 |
| | BRM | -0.001 | 0.000 | 0.005 | 0.005 | 0.009 | 0.151 | 0.039 | 0.089 | 0.956 | 0.872 | 0.940 | 0.853 |
| 7 doses with less extreme values | LRM | 0.000 | -0.000 | 0.000 | 0.001 | 0.005 | 0.030 | 0.016 | 0.026 | 0.891 | 0.828 | 0.883 | 0.799 |
| | BRM | 0.000 | 0.000 | 0.000 | 0.000 | 0.004 | 0.026 | 0.013 | 0.024 | 0.864 | 0.823 | 0.860 | 0.800 |
| 7 doses with extreme values and dose-dependent precision | LRM | -0.016 | -0.047 | 0.137 | 0.111 | 0.069 | 0.409 | 0.583 | 0.520 | 0.965 | 0.573 | 0.921 | 0.685 |
| | BRM | 0.000 | 0.023 | -0.003 | -0.011 | 0.003 | 0.196 | 0.023 | 0.095 | 0.957 | 0.849 | 0.931 | 0.794 |
| **(b) data simulated using normal error term with SD=0.01** | | | | | | | | | | | | | |
| 7 doses with extreme values | LRM | 0.030 | -0.200 | 0.166 | 0.804 | 0.224 | 0.544 | 1.234 | 1.305 | 0.958 | 0.699 | 0.950 | 0.716 |
| | BRM | 0.001 | -0.039 | 0.013 | 0.060 | 0.030 | 0.172 | 0.094 | 0.155 | 0.981 | 0.931 | 0.976 | 0.892 |
| 6 doses after removing largest | LRM | 0.040 | -0.088 | -0.125 | 0.549 | 0.101 | 0.280 | 0.314 | 1.245 | 0.966 | 0.924 | 0.972 | 0.734 |
| | BRM | 0.005 | -0.012 | -0.007 | 0.040 | 0.019 | 0.140 | 0.044 | 0.148 | 0.954 | 0.893 | 0.949 | 0.850 |
| 6 doses after removing smallest | LRM | -0.027 | -0.172 | 0.357 | 0.531 | 0.080 | 0.534 | 0.836 | 1.220 | 0.966 | 0.561 | 0.937 | 0.734 |
| | BRM | -0.005 | -0.035 | 0.025 | 0.040 | 0.018 | 0.178 | 0.078 | 0.143 | 0.958 | 0.877 | 0.943 | 0.857 |
| 7 doses with less extreme values | LRM | 0.000 | -0.001 | 0.001 | 0.004 | 0.011 | 0.060 | 0.032 | 0.053 | 0.892 | 0.826 | 0.883 | 0.799 |
| | BRM | 0.000 | 0.001 | 0.000 | 0.001 | 0.009 | 0.052 | 0.026 | 0.048 | 0.865 | 0.822 | 0.860 | 0.801 |
| 7 doses with extreme values and dose-dependent precision | LRM | -0.052 | -0.215 | 0.524 | 0.462 | 0.130 | 0.622 | 1.123 | 0.984 | 0.965 | 0.521 | 0.921 | 0.726 |
| | BRM | 0.000 | 0.012 | -0.001 | -0.005 | 0.009 | 0.142 | 0.054 | 0.080 | 0.957 | 0.851 | 0.932 | 0.798 |
| **(c) data simulated using normal error term with SD=0.05** | | | | | | | | | | | | | |
| 7 doses with extreme values | LRM | 0.079 | -0.463 | 0.560 | 2.399 | 0.393 | 0.924 | 2.128 | 1.853 | 0.948 | 0.612 | 0.942 | 0.591 |
| | BRM | 0.013 | 0.029 | 0.010 | 0.095 | 0.129 | 0.524 | 0.371 | 0.377 | 0.857 | 0.861 | 0.869 | 0.855 |
| 6 doses after removing largest | LRM | 0.079 | -0.338 | 0.337 | 2.194 | 0.276 | 0.749 | 2.054 | 2.239 | 0.968 | 0.779 | 0.975 | 0.675 |
| | BRM | 0.017 | 0.001 | 0.007 | 0.160 | 0.124 | 0.512 | 0.387 | 0.430 | 0.879 | 0.835 | 0.912 | 0.812 |
| 6 doses after removing smallest | LRM | 0.022 | -0.403 | 0.666 | 2.232 | 0.335 | 0.965 | 1.936 | 2.220 | 0.972 | 0.610 | 0.898 | 0.681 |
| | BRM | 0.009 | -0.021 | 0.030 | 0.160 | 0.119 | 0.511 | 0.358 | 0.420 | 0.874 | 0.754 | 0.862 | 0.819 |
| 7 doses with less extreme values | LRM | 0.005 | -0.077 | 0.086 | 0.362 | 0.084 | 0.383 | 0.693 | 1.215 | 0.905 | 0.795 | 0.895 | 0.813 |
| | BRM | 0.002 | -0.001 | 0.012 | 0.062 | 0.054 | 0.296 | 0.187 | 0.311 | 0.861 | 0.810 | 0.858 | 0.815 |
| 7 doses with extreme values and dose-dependent precision | LRM | -0.029 | -0.441 | 0.994 | 1.679 | 0.297 | 0.955 | 1.840 | 1.715 | 0.945 | 0.444 | 0.904 | 0.678 |
| | BRM | -0.005 | 0.037 | 0.023 | -0.002 | 0.040 | 0.323 | 0.172 | 0.271 | 0.954 | 0.819 | 0.930 | 0.758 |

**Supplemental Table 2.** Simulation result of bias, RMSE and 95% CI coverage probability corresponding to beta error term constant precision

| Scenario | Method | Bias | | | | RMSE | | | | 95% CI Coverage Probability | | | |
|---|---|---|---|---|---|---|---|---|---|---|---|---|---|
| | | $IC_{50}$ | $IC_{90}$ | $\beta_0$ | $\beta_1$ | $IC_{50}$ | $IC_{90}$ | $\beta_0$ | $\beta_1$ | $IC_{50}$ | $IC_{90}$ | $\beta_0$ | $\beta_1$ |
| **(a) data simulated using beta error term with ϕ=35** | | | | | | | | | | | | | |
| 7 doses with extreme values | LRM | 0.017 | -0.304 | 0.142 | 0.653 | 0.163 | 0.481 | 0.658 | 0.686 | 0.924 | 0.697 | 0.909 | 0.566 |
| | BRM | 0.003 | -0.014 | 0.018 | 0.073 | 0.074 | 0.339 | 0.219 | 0.275 | 0.835 | 0.818 | 0.832 | 0.829 |
| 6 doses after removing largest | LRM | 0.039 | -0.174 | -0.027 | 0.504 | 0.115 | 0.402 | 0.354 | 0.666 | 0.933 | 0.864 | 0.936 | 0.666 |
| | BRM | 0.008 | 0.005 | 0.007 | 0.077 | 0.077 | 0.375 | 0.229 | 0.298 | 0.812 | 0.808 | 0.811 | 0.803 |
| 6 doses after removing smallest | LRM | -0.022 | -0.288 | 0.248 | 0.502 | 0.108 | 0.501 | 0.499 | 0.670 | 0.930 | 0.620 | 0.901 | 0.664 |
| | BRM | -0.002 | -0.020 | 0.031 | 0.074 | 0.075 | 0.353 | 0.223 | 0.298 | 0.815 | 0.782 | 0.805 | 0.800 |
| 7 doses with less extreme values | LRM | 0.004 | -0.045 | 0.023 | 0.117 | 0.063 | 0.332 | 0.195 | 0.300 | 0.900 | 0.828 | 0.891 | 0.821 |
| | BRM | 0.003 | 0.031 | 0.004 | 0.028 | 0.058 | 0.331 | 0.170 | 0.272 | 0.845 | 0.837 | 0.847 | 0.838 |
| 7 doses with extreme values and dose-dependent precision | LRM | 0.080 | -0.093 | -0.179 | 0.444 | 0.147 | 0.281 | 0.435 | 0.548 | 0.904 | 0.908 | 0.927 | 0.639 |
| | BRM | 0.003 | 0.031 | -0.003 | 0.012 | 0.067 | 0.277 | 0.182 | 0.236 | 0.755 | 0.780 | 0.768 | 0.772 |
| **(b) data simulated using beta error term with ϕ=15** | | | | | | | | | | | | | |
| 7 doses with extreme values | LRM | 0.029 | -0.571 | 0.373 | 1.633 | 0.233 | 0.555 | 1.177 | 1.197 | 0.941 | 0.588 | 0.910 | 0.406 |
| | BRM | 0.008 | -0.050 | 0.036 | 0.174 | 0.109 | 0.464 | 0.324 | 0.403 | 0.817 | 0.790 | 0.820 | 0.805 |
| 6 doses after removing largest | LRM | 0.074 | -0.360 | -0.043 | 1.269 | 0.164 | 0.516 | 0.635 | 1.192 | 0.942 | 0.823 | 0.939 | 0.556 |
| | BRM | 0.018 | -0.028 | 0.019 | 0.203 | 0.117 | 0.527 | 0.355 | 0.446 | 0.778 | 0.783 | 0.794 | 0.776 |
| 6 doses after removing smallest | LRM | -0.042 | -0.564 | 0.620 | 1.280 | 0.157 | 0.596 | 0.890 | 1.180 | 0.939 | 0.503 | 0.905 | 0.555 |
| | BRM | -0.001 | -0.071 | 0.066 | 0.192 | 0.115 | 0.499 | 0.345 | 0.449 | 0.787 | 0.733 | 0.782 | 0.777 |
| 7 doses with less extreme values | LRM | 0.008 | -0.104 | 0.059 | 0.286 | 0.096 | 0.513 | 0.320 | 0.489 | 0.906 | 0.795 | 0.884 | 0.807 |
| | BRM | 0.006 | 0.074 | 0.012 | 0.062 | 0.088 | 0.553 | 0.260 | 0.409 | 0.838 | 0.836 | 0.841 | 0.847 |
| 7 doses with extreme values and dose-dependent precision | LRM | 0.165 | -0.202 | -0.425 | 1.114 | 0.218 | 0.374 | 0.771 | 0.959 | 0.916 | 0.886 | 0.941 | 0.534 |
| | BRM | 0.010 | 0.053 | -0.008 | 0.050 | 0.104 | 0.423 | 0.281 | 0.368 | 0.763 | 0.790 | 0.764 | 0.769 |
| **(c) data simulated using beta error term with ϕ=5** | | | | | | | | | | | | | |
| 7 doses with extreme values | LRM | 0.042 | -0.851 | 0.748 | 3.414 | 0.270 | 0.527 | 1.611 | 1.421 | 0.948 | 0.423 | 0.898 | 0.198 |
| | BRM | 0.008 | -0.011 | 0.047 | 0.191 | 0.161 | 0.606 | 0.589 | 0.767 | 0.811 | 0.800 | 0.815 | 0.811 |
| 6 doses after removing largest | LRM | 0.106 | -0.636 | 0.181 | 2.962 | 0.234 | 0.865 | 1.401 | 1.604 | 0.949 | 0.684 | 0.938 | 0.356 |
| | BRM | 0.007 | -0.115 | 0.112 | 0.355 | 0.166 | 0.643 | 0.621 | 0.761 | 0.762 | 0.752 | 0.773 | 0.769 |
| 6 doses after removing smallest | LRM | -0.039 | -0.871 | 1.126 | 2.935 | 0.231 | 0.649 | 1.428 | 1.620 | 0.953 | 0.380 | 0.883 | 0.362 |
| | BRM | 0.008 | -0.087 | 0.070 | 0.328 | 0.166 | 0.588 | 0.509 | 0.724 | 0.766 | 0.755 | 0.772 | 0.773 |
| 7 doses with less extreme values | LRM | 0.008 | -0.456 | 0.275 | 1.126 | 0.152 | 0.515 | 0.679 | 1.041 | 0.915 | 0.639 | 0.874 | 0.707 |
| | BRM | 0.002 | -0.088 | 0.093 | 0.318 | 0.132 | 0.588 | 0.423 | 0.660 | 0.839 | 0.812 | 0.834 | 0.835 |
| 7 doses with extreme values and dose-dependent precision | LRM | 0.245 | -0.378 | -0.646 | 2.300 | 0.274 | 0.470 | 1.207 | 1.262 | 0.908 | 0.833 | 0.947 | 0.343 |
| | BRM | 0.025 | 0.171 | -0.032 | 0.074 | 0.161 | 0.762 | 0.426 | 0.549 | 0.757 | 0.796 | 0.761 | 0.770 |

**Supplementary Table 3.** The first example of REAP application with B-cell lymphoma data, corresponding with Figure 5. $IC_{50}$ estimations are ranked from low to high. Hypothesis testings on equal potency (i.e., concentration for $IC_{50}$) were conducted pairwise with the group right above (one rank lower). Jeko-1 has the highest potency and the difference of $IC_{50}$ estimations between Jeko-1 and Jeko-R is significant with a p-value < 0.0001. The B-cell lymphoma dataset is available at https://github.com/vivid225/REAP/blob/main/REAP/.

| Model | Intercept | Slope (m) | Std. Err for m | P-value for m>1 | $IC_{50}$ estimation | Std. Err for $IC_{50}$ estimation | Pairwise comparison |
|---|---|---|---|---|---|---|---|
| Jeko-1 | -4.807 | -1.252 | 0.155 | 0.0519 | 0.021 | 0.008 | - |
| Jeko-R | -4.305 | -1.822 | 0.112 | <.0001 | 0.094 | 0.006 | <0.0001 |
| Rec-1 | -5.304 | -2.63 | 0.091 | <.0001 | 0.133 | 0.004 | <0.0001 |
| Mino | -2.684 | -1.474 | 0.141 | 0.0004 | 0.162 | 0.015 | 0.0656 |
| Jeko-NO #1 | -2.012 | -1.192 | 0.135 | 0.0769 | 0.185 | 0.021 | 0.3755 |
| MAVER-1 | -2.21 | -1.37 | 0.125 | 0.0015 | 0.199 | 0.021 | 0.6312 |
| Jeko-NO #11 | -1.459 | -1.267 | 0.152 | 0.0398 | 0.316 | 0.038 | 0.0114 |
| JVM2 | -1.056 | -1.271 | 0.135 | 0.0223 | 0.436 | 0.055 | 0.0818 |

**Supplementary Table 4.** The output for the estimated dose-response curve of anti-viral drugs under the same biological batch with SARS-CoV-2 data. Calpain inhibitor IV has the highest potency (p-value = 0.0038). The reconstructed SARS-CoV-2 dataset is available at https://github.com/vivid225/REAP/blob/main/REAP/

| Model | Intercept | Slope (m) | Std. Err for m | P-value for m>1 | $EC_{50}$ estimation | Std. Err for $EC_{50}$ estimation | Pairwise comparison |
|---|---|---|---|---|---|---|---|
| CalpainInhibitorIV | 0.678 | 0.725 | 0.114 | 0.9918 | 0.393 | 0.103 | - |
| Chloroquine | -1.013 | 0.84 | 0.135 | 0.8813 | 3.337 | 0.88 | 0.0038 |
| Remdesivir | -1.791 | 0.797 | 0.12 | 0.9553 | 9.469 | 2.638 | 0.0282 |
| Hydroxychloroquine | -1.485 | 0.562 | 0.075 | 1 | 14.074 | 4.994 | 0.4445 |
| E64d (Aloxistatin) | -3.211 | 0.861 | 0.129 | 0.8587 | 41.61 | 15.473 | 0.1242 |
